# The prognostic effects of somatic mutations in ER-positive breast cancer

**DOI:** 10.1101/235846

**Authors:** Obi L Griffith, Nicholas C Spies, Meenakshi Anurag, Malachi Griffith, Jingqin Luo, Dongsheng Tu, Belinda Yeo, Jason Kunisaki, Christopher A Miller, Kilannin Krysiak, Jasreet Hundal, Benjamin J Ainscough, Zachary L Skidmore, Katie Campbell, Runjun Kumar, Catrina Fronick, Lisa Cook, Jacqueline E Snider, Sherri Davies, Shyam M Kavuri, Eric C Chang, Vincent Magrini, David E Larson, Robert S Fulton, Shuzhen Liu, Samuel Leung, David Voduc, Ron Bose, Mitch Dowsett FMedSci, Richard K Wilson, Torsten O Nielsen, Elaine R Mardis, Matthew J Ellis

## Abstract

More than 50 genes are recurrently affected by somatic mutation in estrogen receptor positive (ER+) breast cancer but prognostic effects have not been definitively established. Primary tumor DNA was therefore subjected to targeted sequencing from 625 postmenopausal (UBC-TAM series) and 328 premenopausal (MA12 trial) hormone receptor-positive (HR+) patients. Independent validation of prognostic interactions was achieved using independent data from the METABRIC study. Associations between MAP3K1 and PIK3CA with luminal A status and TP53 mutations with Luminal B/non-luminal tumors were observed, validating the methodological approach. In UBC-TAM, *NF1* frame-shift nonsense (*FS/NS*) mutation was validated as a poor outcome driver. For MA12, poor outcome associated with PIK3R1 mutation was similarly validated. DDR1 mutations were strongly associated with poor prognosis in UBC-TAM despite stringent false-discovery correction (q=0.0003). In conclusion, uncommon recurrent somatic mutations should be further explored to create a more complete explanation of the highly variable outcomes that typify ER+ breast cancer.

## Introduction

While recent genomic studies have provided a comprehensive catalog of genes that accumulate somatic point mutations and small insertions/deletions (indels) in estrogen receptor-positive (ER+) breast cancer, there remains considerable uncertainty as to how these newly discovered mutations relate to disease outcomes^1–3^. Most genomic discovery cohorts were neither uniformly treated nor followed long enough. For ER+ disease in particular, prognostic studies require prolonged observation since late relapses can occur^4^. Uniform treatment was a feature of a whole genome sequencing study of samples accrued from a neoadjuvant aromatase inhibitor (AI) clinical trial for ER+ clinical stage 2 or 3 disease, although only short-term anti-proliferative response to AI was reported. This investigation identified that mutations in *MAP3K1*, a tumor suppressor gene involved in stress kinase activation, were associated with indolent biological features and low proliferation rates^5^. The resulting hypothesis was that *MAP3K1* mutation would be associated with favorable outcomes. In contrast, *TP53* mutations associated with poor prognosis features and high proliferation rates.

To more comprehensively address the relationships between somatic mutations and outcomes in ER+ breast cancer, we developed an approach to detect somatic mutations in DNA isolated from formalin fixed tumor blocks that were over 20 years old. After curating existing mutational data from breast cancer genomics discovery studies (Supplementary Data 1), 83 genes were chosen for analysis (Supplementary Table 1). We applied DNA hybrid capture, sequencing and somatic analysis to three ER+ breast cancer discovery cohorts with contrasting clinical characteristics: An older cohort treated with adjuvant tamoxifen and no chemotherapy, a premenopausal cohort uniformly treated with chemotherapy and randomized to tamoxifen versus observation; and a third mixed cohort that was used to expand the mutational landscape analysis (Supplementary Table 2). An analytical pipeline was developed to identify somatic variants while compensating for the lack of matched normal DNA, which is generally unavailable in the setting of older formalin-fixed tumor material. Somatic mutations were analyzed for association with standard clinical variables, wherein mutated *TP53* and *MAP3K1* served as *a priori* hypotheses for poor and good outcome, respectively. Additional objectives were to identify new mutational hotspots and to determine mutation frequencies for therapeutic targets. Validation was possible by comparing our results to those in cBioPortal where the mutational analysis in the METABRIC cohort overlapped with the 83 genes investigated in the study described here.

## Results

### Sequencing and final study cohorts

University of British Columbia Tamoxifen Series (UBC-TAM): These cases were drawn from a well-annotated cohort of patients treated with adjuvant tamoxifen without chemotherapy^6^. A total of 625 of 632 (98.8%) patient samples that fully met study criteria passed a minimum sequencing quality cutoff of at least 80% of targeted bases covered at greater than 20X (mean coverage: 133X) with other quality metrics described in the supplementary data (Supplementary Figure 1-5 and Supplementary Data 2). The final patient population had an average age of 67 at diagnosis (range: 40-89+). All were treated with five years of adjuvant tamoxifen, and were primarily postmenopausal, grade 2 or 3 cancers, of ductal histologic subtype (Supplementary Table 2). All were ER+ and at least 88.6% were clinically HER2- (13/625 unknown). A subset of 463 of these patients had PAM50 subtyping data available from a previous study ^6^. The median follow up in the cohort examined was 25 years and one month.

POLAR cohort: This patient series was a case-control study of ER+ breast tumors, 175 of 194 (90.2%) patient samples passed minimum sequencing quality thresholds. A case was defined as any patient who relapsed during follow-up, and controls were defined as lacking relapse through a similar follow-up duration. Based on these definitions, there were 91 cases and 84 controls. Of the cases, 43 were early relapses (<5 years since diagnosis) and 48 were late relapses (>5 years). Patients were only included if they received adjuvant endocrine therapy, but chemotherapy was not an exclusion criterion, nor was menopausal status. These cases were used in the mutation landscape and hotspot analyses only.

NCIC-MA12 Trial cohort. These cases were drawn from a clinical trial in premenopausal women treated with a standard adjuvant chemotherapy regimen and randomized to tamoxifen versus observation. A total of 459 patient samples passed the minimum sequencing quality threshold, of which 328 were hormone receptor positive (HR+), and only the HR+ cohort are included here for most analyses. The majority were premenopausal (mean age of 45). All patients received chemotherapy, and 48% were treated with 5 years of adjuvant tamoxifen. A subset of 255 of these patients had PAM50 subtyping data available. The median follow up in the cohort examined was 9.7 years

Across the three cohorts, there were 1,259 patient samples that passed minimum sequencing quality thresholds and 1,128 of these were ER+ (UBC-TAM and POLAR) or HR+ (MA12).

### Variant calling and filtering

A total of over 62 million variants were called in UBC-TAM. After extensive filtering against a set of nearly 70,000 unmatched normal samples and manual review to eliminate common polymorphisms and false positives (see methods), 1,991 putative somatic variants were identified (0 to 26 variants per patient). A set of 1,693 mutations was defined as the “non-silent” set for further analysis that excluded sequencing variants in splice regions, RNA genes (except *MALAT1*), UTRs, introns, and all silent mutations. Finally, a set of 408 frameshift or nonsense mutations was defined. The same filtering method was applied to both the POLAR and MA12 datasets. A total of 540 putative somatic mutations (436 non-silent, 145 FS/NS) were identified in POLAR, and 2,104 (1,753 non-silent, 610 FS/NS) in MA12. Full details on these variants are included in Supplementary Data 3 and summarized for key genes in Supplementary Figure 6.

### Mutation landscape analysis

In 1128 samples passing quality control standards, considering only non-silent mutations, 17 genes were mutated at a rate greater than 5%, and 6 at a rate greater than 10%; *PIK3CA* was the only gene mutated in greater than 20% of samples (**Figure 1A**). The order from most recurrent to least for the 10 most frequently mutated genes was: *PIK3CA* (41.1%), *TP53* (15.5%), *MLL3* (13.4%), *MAP3K1* (12.0%), CDH1 (10.5%), MALAT1 (10.0%), GATA3 (9.1%), MLL2 (8.7%), ARID1A (7.2%), and BRCA2 (6.6%). This list correlates well with previously reported recurrently mutated genes. For example, the top 4 most significantly mutated genes in the ER+ subset of TCGA breast proiect^3^ were *PIK3CA* (24.3%), *TP53* (14.6%), *GATA3* (8.9%) and *MAP3K1* (6.2%). The overall average mutation rate was estimated as 3.3 per MB of coding sequence (range: 0.5 to 13.8 mutations per MB, excluding samples with no mutations called). In order to determine whether mutations in any gene pair were mutually exclusive or co-occuring in this dataset, a pairwise Chi-squared or Fisher’s exact test was performed. Mutations in PIK3CA and MAP3K1 were significantly more likely to co-occur (after BH FDR correction) in TAM dataset, and were near significance in MA12 although not after correction (p = 0.08). These results are summarized in Supplementary Data 4.

**Figure 1.**
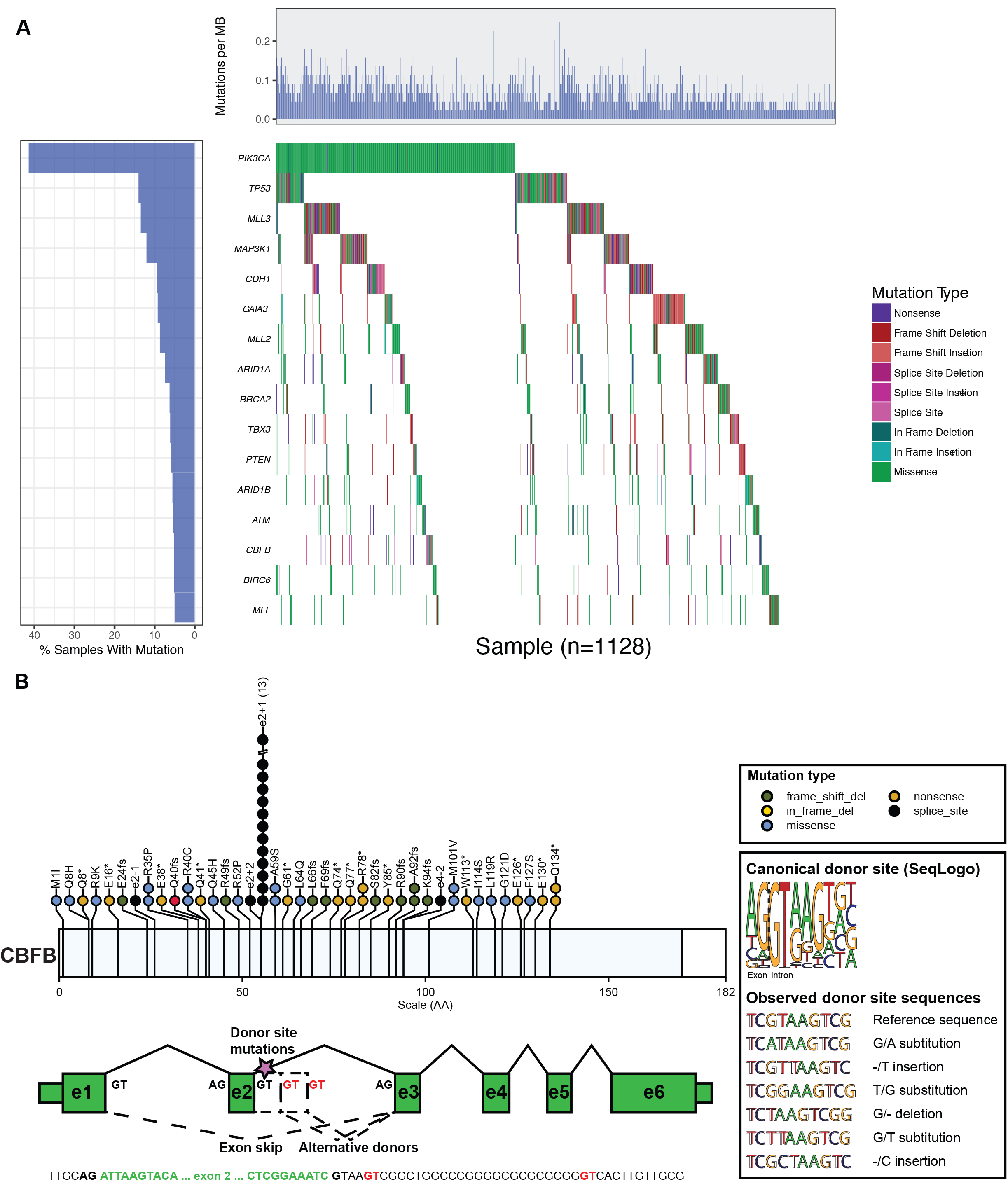
Mutation recurrence and novel splice site mutation. A) The overall mutation recurrence rate ranged from 41.1% of samples for *PIK3CA* to 0.0% for *PIN1*. The figure depicts non-silent mutations for all 1128 patients for the top 16 most recurrently mutated genes (>5% recurrence). If a patient had multiple mutations it is colored according to the “most damaging” mutation following the order presented in the Mutation Type legend (vertical color bar). Mutations per MB were calculated using the total number of mutations observed over the total exome space corresponding to the tiled space from “SeqCap EZ Human Exome Library v2.0”. A correction factor was applied for genes not assayed using the expected number of additional mutations based on TCGA data. B) Mutation recurrence rates (amino acid level) in this study were compared to previously reported mutation rates from a multi-study MAF file of six reported breast cancer sequencing studies (Supplementary Data 1). An entirely novel mutation “hot spot” was discovered affecting the exon 2 splice (donor) site of *CBFB* in at least 15 patients. Six different single nucleotide substitutions, insertions and deletions were observed, all affecting either the first or second base of the donor splice site. These mutations were most likely missed in previous studies because of a lack of sequencing coverage due to the GC-rich nature of exons 1 and 2 of *CBFB* (Supplementary Figures 9-10). Such mutations are predicted to significantly alter the canonical donor site and result in either alternate donor usage or skipping of one or more exons of *CBFB*.

### Hotspot analysis

As anticipated^7^, mutations in *PIK3CA* at *E542K, E545K*, and *H1047R* were highly recurrent in this study with 69/1259 (5.5%) E542K, 104 (8.3%) E545K, and 181 (14.4%) H1047R mutations (Supplementary Figure 6C). Mutations in the ligand binding domain of *ESR1* (1.1%) were extremely rare^3^ (Supplementary Figure 6A). To uncover novel hotspots in these data, both Chi-squared and Fisher’s exact tests were performed using mutation frequencies from previous sequencing studies as the expected values (see Methods for definition of multi-study MAF file) (Supplementary Table 3). The most notable novel finding was in *CBFB* (**Figure 1B**). At least 6 different genomic alterations were observed in 15 patients (Supplementary Data 3) that affected the donor splice site of exon 2. Manual review of this splice site identified at least two additional patients with evidence for mutations at this location. The predicted effect of these mutations is skipping of exon 2 or alternate donor site usage, each likely resulting in loss-of-function of the *CBFB* protein. Additional splice site mutations were observed at the exon 2, exon 4 and exon 5 acceptor sites of *CBFB*. ErbB2 expressed the anticipated profile of activating mutations from earlier publications^8^ with 22/1259 (1.7%) samples harboring known activating mutations and another 6 variants of unknown significance in the kinase domain or at the S310 residue (**Figure 8C**).

### Somatic mutation association with PAM50-based intrinsic subtype

The PAM50 intrinsic subtype calls were obtained from previously published analyses to compare their mutational profiles between UBC-TAM and the MA12 studies. In both studies about half the patients had luminal A tumor. However, the MA12 cohort had a higher proportion of non-luminal subtypes, with 19.8% HER2-E and 6.6% basal and fewer luminal B tumors (25.1% versus 42.4%) (**Figure 2A-B**). Age density plots by subtype serve to emphasize the large difference in the median age between the two sample cohorts (43 versus 65), and also the influence of age with respect to the intrinsic subtype incidence. Namely, in the younger MA12 cohort, there is a younger peak incidence with basal-like breast cancer than Luminal A disease (**Figure 2D**). In contrast in the older UBC-TAM cohort, an influence of age on intrinsic subtype was not observed (**Figure 2C**). Relationships between intrinsic subtype and mutation patterns were also explored, classifying mutation positive status as “non-silent”, “missense”, nonsense/frame-shift (FS/NS) or FS/NS+splice site (Supplementary Data 5). The FDR corrected p-value (q-value) took into account that 83 genes were examined. However, this level of false discovery detection could be viewed as overly conservative in an exploratory analysis and any gene mutation with q-value association of <0.2 was therefore considered reportable^9–11^. For MA12, non-silent TP53 mutation was highly subtype-associated because of the very high incidence in non-luminal versus luminal subtypes. PIK3CA and MAP3K1 mutations were associated with Luminal A disease in both cohorts (Supplementary Figure 7A). Finally, there was a strong association between Luminal B status and non-silent (Supplementary Figure 7B) as well as FS/NS mutations in GATA3 (Supplementary Data 5, q value = 0.006). GATA3 mutations were present in 28-30% of Luminal B cases and less so in luminal A cases (5%). Considering q values of <0.2 the associations between FS/NS and non-silent mutations in ATM and Luminal B tumors in MA12 (8-13%) suggests that ATM loss is also a possible luminal B driver (Supplementary Figure 7B), at least in younger women (MA12). Relationships between age and mutation incidence were therefore also explored (Supplementary Figure 7C), with the finding that both ATM mutation and GATA3 mutations were associated with an earlier age of onset within the luminal B category (**Figure 2E and 2F**). Finally, NF1 mutations were associated with the HER2-enriched subtype in the UBC-Tam series, explaining the association with poor outcomes (Supplementary Figure 7B).

**Figure 2.**
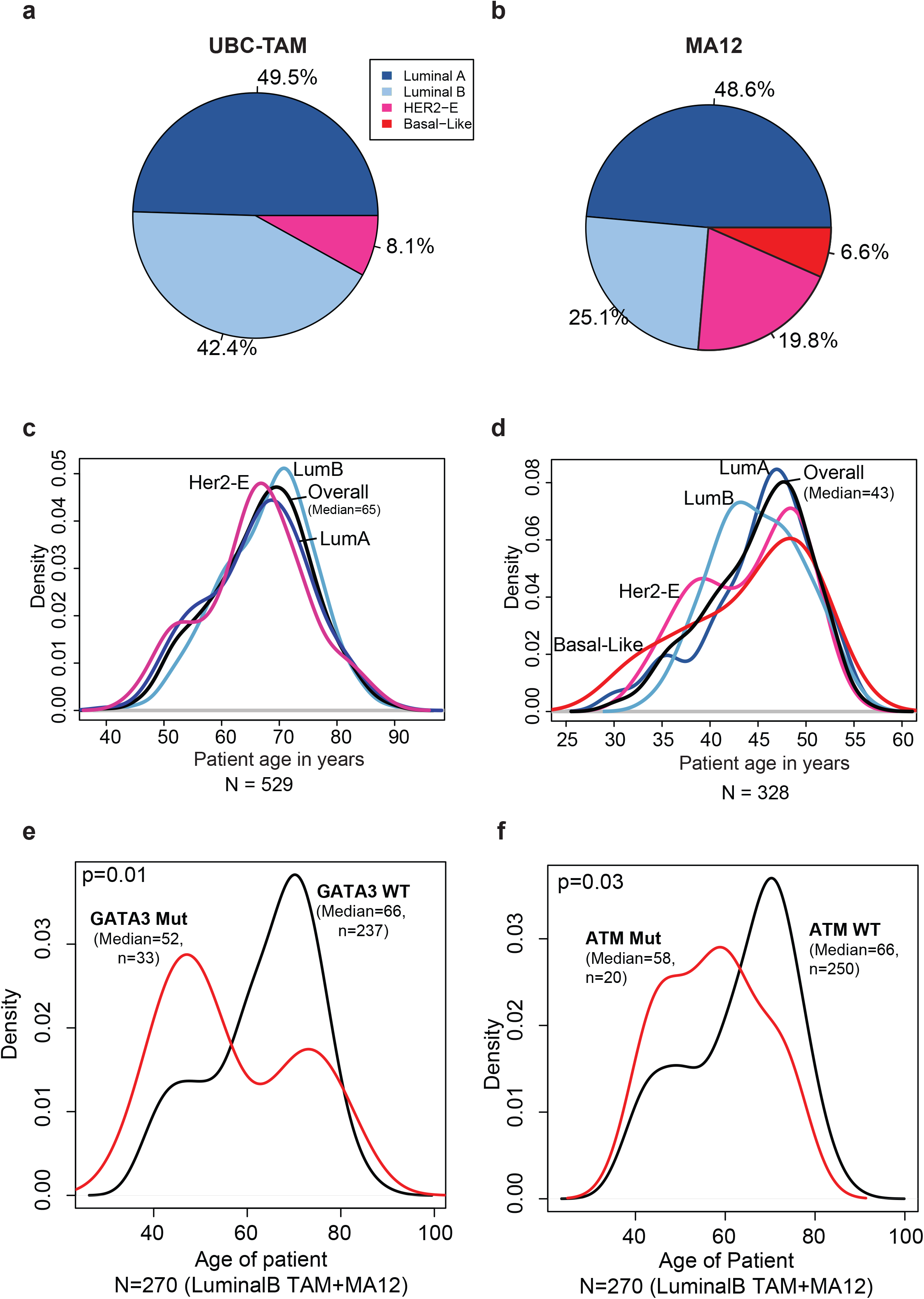
Cross-cohort age and subtype analysis. A-B) Percentage composition of samples by intrinsic subtype of the tumor in the two discovery cohorts for UBC-TAM (A) and MA12 (B) cohorts. C-D) Age-density plots for patients categorized by intrinsic subtype in UBC-TAM (C) and MA12 (D) cohorts. The overall median age shows that UBC-TAM is constituted mostly of post-menopausal patients (median age=65), in contrast to MA12, which has younger patients (median age=43). E-F) Younger luminal B subtype patients harbor GATA3 (E) and ATM (F) mutations in the combined set of UBC-TAM and MA12 Luminal B cases (median age=52, p=0.01; median age=58, p=0.03 for GATA3 and ATM respectively).

### Survival analysis according to somatic mutation

For the UBC-TAM Series (**Figure 3A**), univariate analysis of favorable prognostic associations for breast cancer specific survival (BCSS) were detected for non-silent mutations in *MAP3K1, ERBB3*, XBP1 and PIK3CA (**Figure 3B**, Supplementary Data 6). Adverse prognostic effects were observed for non-silent mutations in *DDR1* and *TP53*, as well as for frame-shift and nonsense (FS/NS) mutations in NF1. An analysis for recurrence free survival (RFS) produced similar results, except for ARID1B, which was marginally associated with more favorable outcome. A multivariate Cox model was applied to put each gene in the context of clinical parameters (grade, tumor size and node status). These analyses indicated that the prognostic effects of non-silent DDR1, PIK3CA, GATA3 FS/NS, TP53 and MAP3K1 mutations were independent of grade and pathological stage (**Figure 3C**). Multiple correction testing, yielded DDR1 as the only gene that remained significant with a q-value of 0.0003. (Supplementary Data 5). For the MA12 clinical trial cohort (**Figure 4A**) we focused on overall survival associations as this was the primary endpoint of the study and the most robust endpoint. A number of rarely mutated genes were associated with poor outcome in univariate analysis as displayed in **Figure 4B**. Multiple testing corrections indicated none of these findings could be considered significant^9–11^. However, in multivariate analysis, based on the uncorrected p value, the prognostic effects of mutations in ErbB2, ErbB4, LTK FS/NS, MAP3K4, PIK3R1, RB1, RELN and TGFB2 were independent of pathological stage and grade (**Figure 4B**).

**Figure 3.**
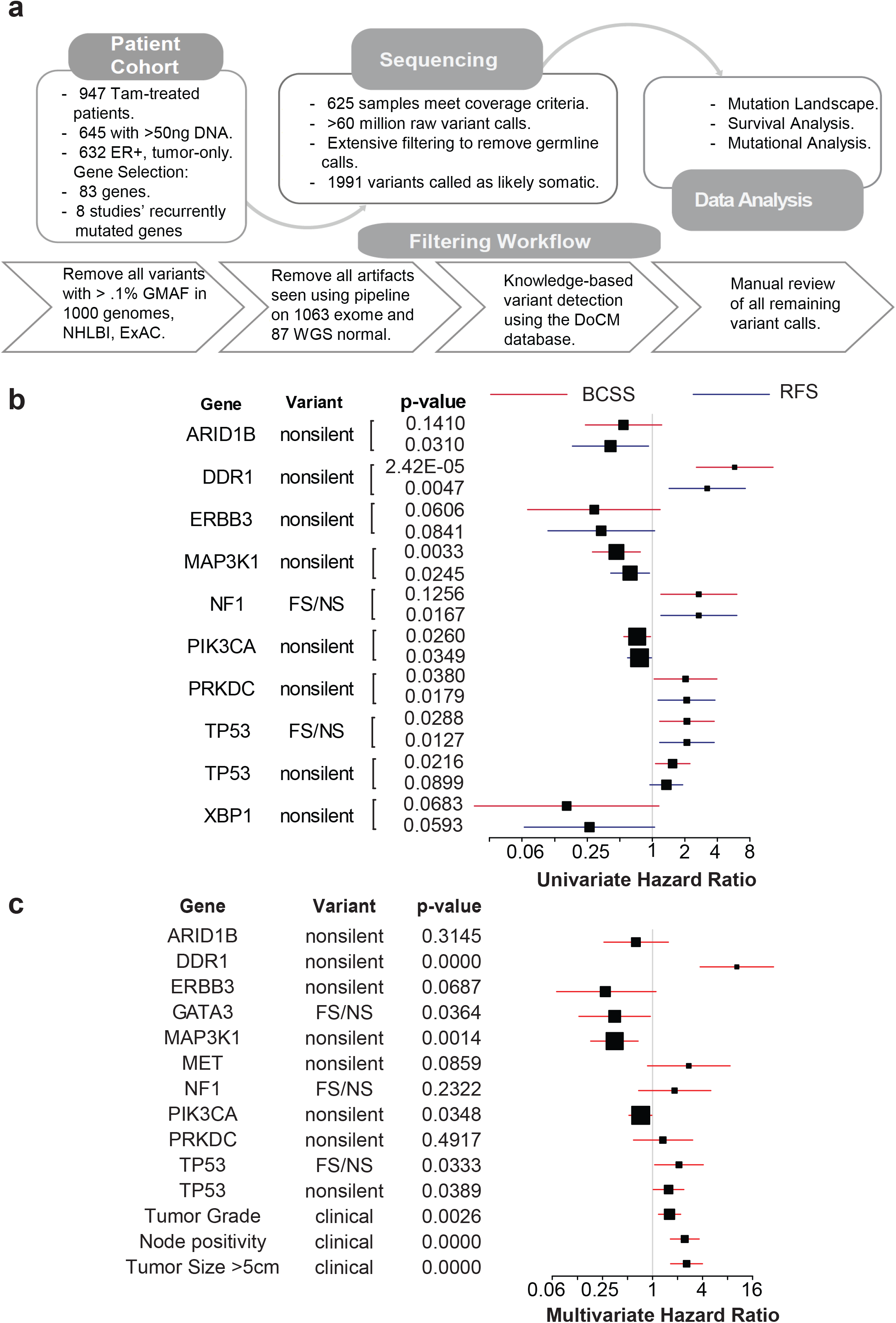
Candidate discovery from UBC-TAM cohort and prognosis evaluation. (A) DNA was extracted from tumor specimens from 947 patients with ER+ breast cancer treated with tamoxifen monotherapy for 5 years. 632 samples with adequate yield were sequenced for 83 genes known to be recurrently mutated or breast cancer relevant. A total of 625 samples passed minimum quality checks and were sequenced to an average of 135.8X coverage. A total of ~62 million variants from the reference genome were identified. Extensive filtering and manual review reduced this list to 1,991 putatively somatic variants. Survival analysis was applied to non-silent and truncating gene mutation status versus disease outcome (relapse or breast-cancer-specific death). In addition, mutations were analyzed for novel hotspots, patterns of mutual exclusivity or co-occurrence and association with clinical variables. (B) Forest plot of impact of mutations in candidate genes, identified using UBC-TAM population, on breast-cancer-specific-survival (red) and recurrence-free survival (blue). The variant types are characterized based on non-silent or nonsense/frameshift (FS/NS) mutations. The box size is relative to frequency of mutations listed in the analysis, larger boxes represent high incidence rate mutations. (C) Multivariate forest plot of effect of mutations in UBC-TAM candidate genes on breast cancer specific-survival when assessed together with clinical factors including Tumor Grade, Node positivity and Tumor Size (>5cm).

**Figure 4.**
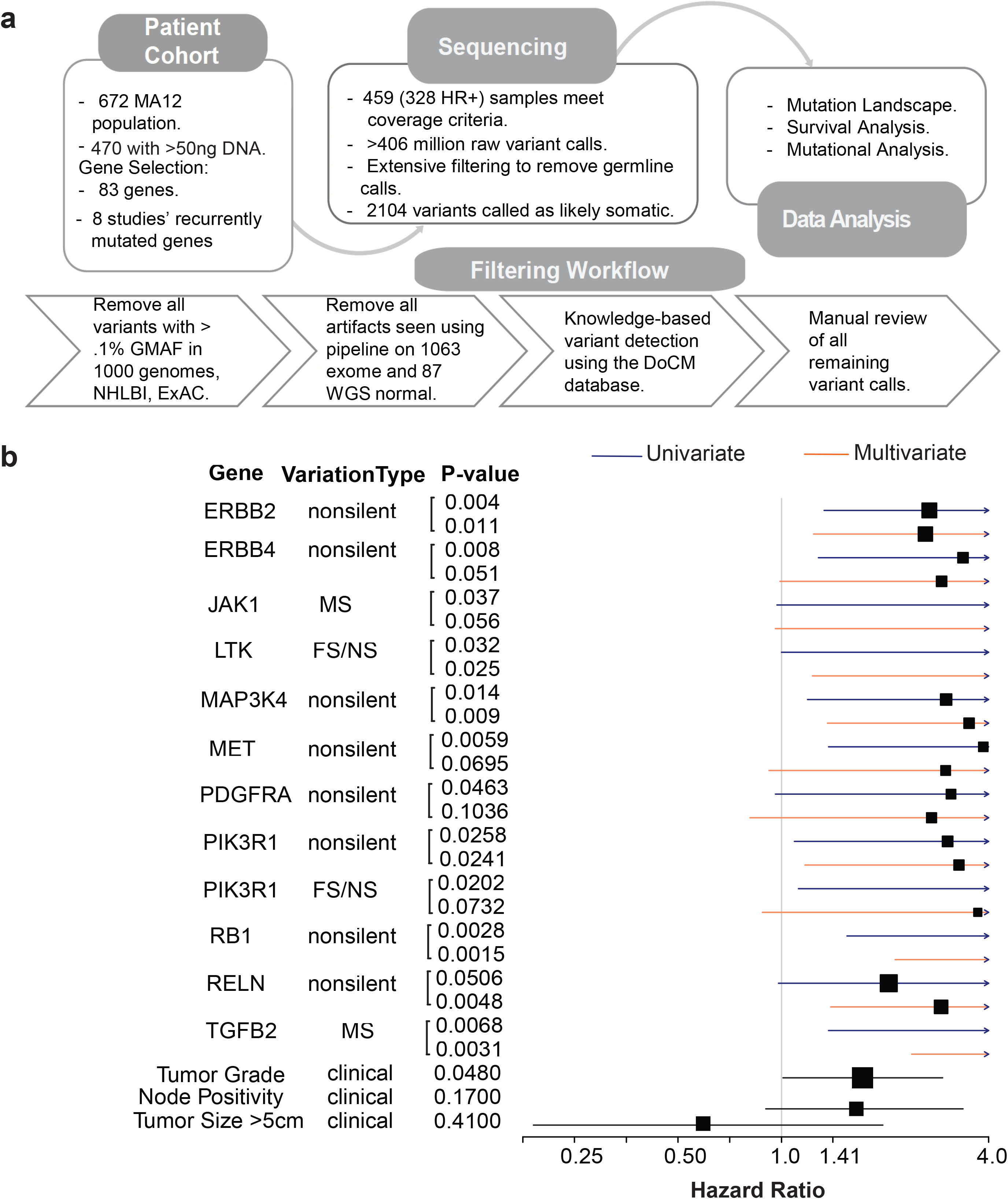
Candidate discovery from MA12 cohort and prognosis evaluation. (A) DNA was extracted from tumor specimens and 470 samples with adequate yield were sequenced for 83 genes known to be recurrently mutated or breast cancer relevant. A total of 459 (328 HR+) samples passed minimum quality checks and were sequenced to an average of 272.6X coverage. A total of 406 million variants from the reference genome were identified. Extensive filtering and manual review reduced this list to 2104 putatively somatic variants. Survival analysis was applied to non-silent and truncating gene mutation status versus overall survival. (B) Forest plot showing effect of mutation in candidate genes on overall survival (univariate - blue, multivariate - orange), along with the clinical factors used in the multivariate analysis, tumor grade, node positivity and tumor size (>5cm) in black. The box size is relative to frequency of mutations listed in the analysis, larger boxes represents high incidence rate mutations. Note: a few boxes are not shown if their hazard ratio were greater than 4.0.

### Verification of Prognostic effects of Mutations in METABRIC data

While few genes were significant in univariate analysis after multiple testing correction, they provide valuable hypotheses for further testing and validation. We therefore sought additional data in the public domain to further assess the uncorrected p value-based findings in our data set. The METABRIC consortium have reported somatic mutations in cBioPortal^12^ with co-reported detailed hormone receptor status, age at diagnosis (median age=64 years for ER+ patients), mean follow up of >8 years and disease-specific outcome^13, 14^. This data set provided the opportunity to conduct a validation exercise for overlapping genes in the two data sets. For the UBC-TAM series (**Figure 3**), 9 genes with a univariate p value of <0.05 were brought forward for validation (**Figure 5**). Of the 6 overlapping genes also examined in METABRIC, consistent prognostic effects independent of clinical variables were observed for non-silent mutations in three genes, *MAP3K1* (favorable), *TP53* (unfavorable) and *NF1* FS/NS mutations (unfavorable). For the MA12 series (**Figure 4**), 5 shared genes were identified with univariate p values of <0.05, yet only *PIK3R1* mutations (non-silent or FS/NS) showed consistent adverse prognostic effects (**Figure 6**). The Kaplan Meier survival plots for the consistent adverse prognostic effects of *NF1* FS/NS and non-silent *PIK3R1* mutations are illustrated in **Figure 7A-D**.

**Figure 5.**
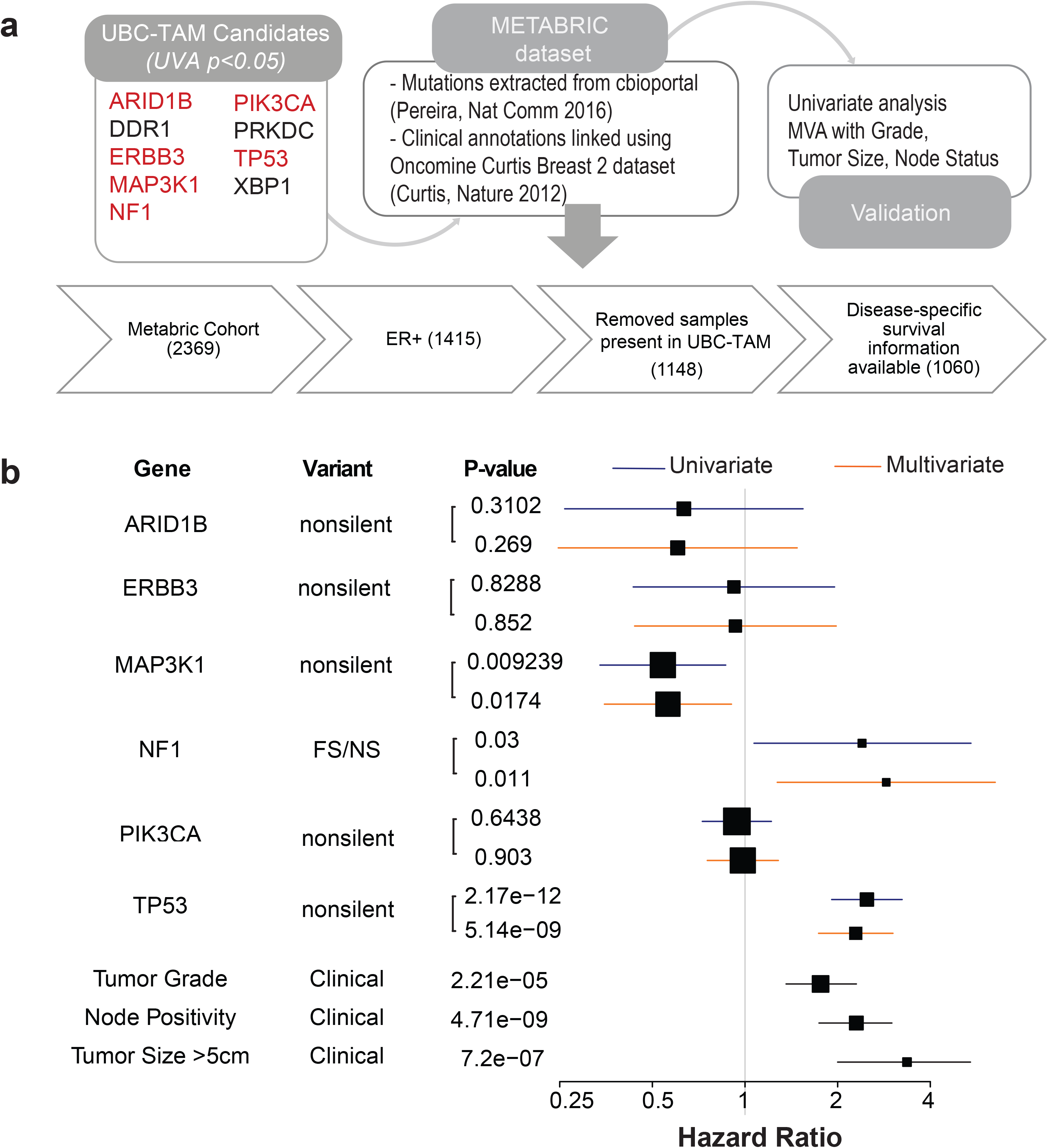
Validation of UBC-TAM candidates in ER+ METABRIC. A) Six out of nine candidate genes from UBC-TAM analysis had mutations reported in METABRIC cohort. 1060 ER+ samples with disease-specific survival information were used to test the effect of mutations in the candidate genes on prognosis. B) Forest plot shows effect of mutated candidate genes on disease-specific survival in METABRIC ER+ cohort with univariate cox proportional-hazard ratio in blue and multivariate in orange. The clinical factors used in the multivariate analysis, namely tumor grade, node positivity and tumor size (>5cm), are shown in black. The box size is relative to frequency of mutations listed in the analysis, larger boxes represent genes with higher incidence rate of mutations.

**Figure 6.**
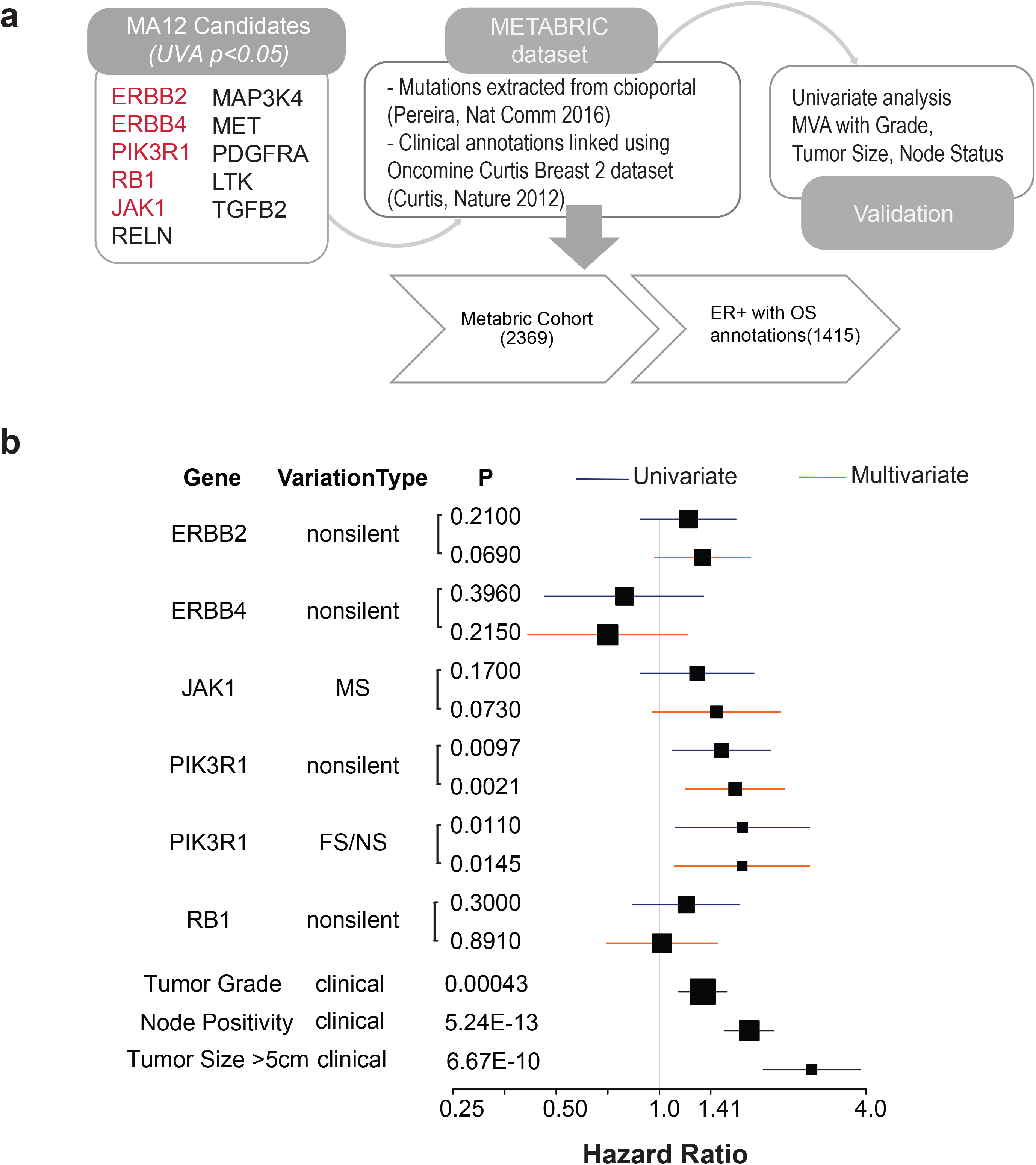
Validation of MA12 candidates in ER+ METABRIC. A) Five out of eleven candidates from MA12 analysis had mutations reported in the METABRIC cohort. 1415 ER+ samples with overall survival information was used to test the effect of mutations in the candidate genes on prognosis. B) Forest plot shows effect of mutated candidate genes, shortlisted based on MA12 mutation analysis, on overall survival in METABRIC ER+ breast cancer patients. Univariate (blue) and multivariate (orange) cox proportional-hazard ratio depict the independent prediction of survival outcomes for the six candidate genes. The box size is relative to frequency of mutations listed in the analysis, larger boxes represent genes with higher incidence rate of mutations.

**Figure 7.**
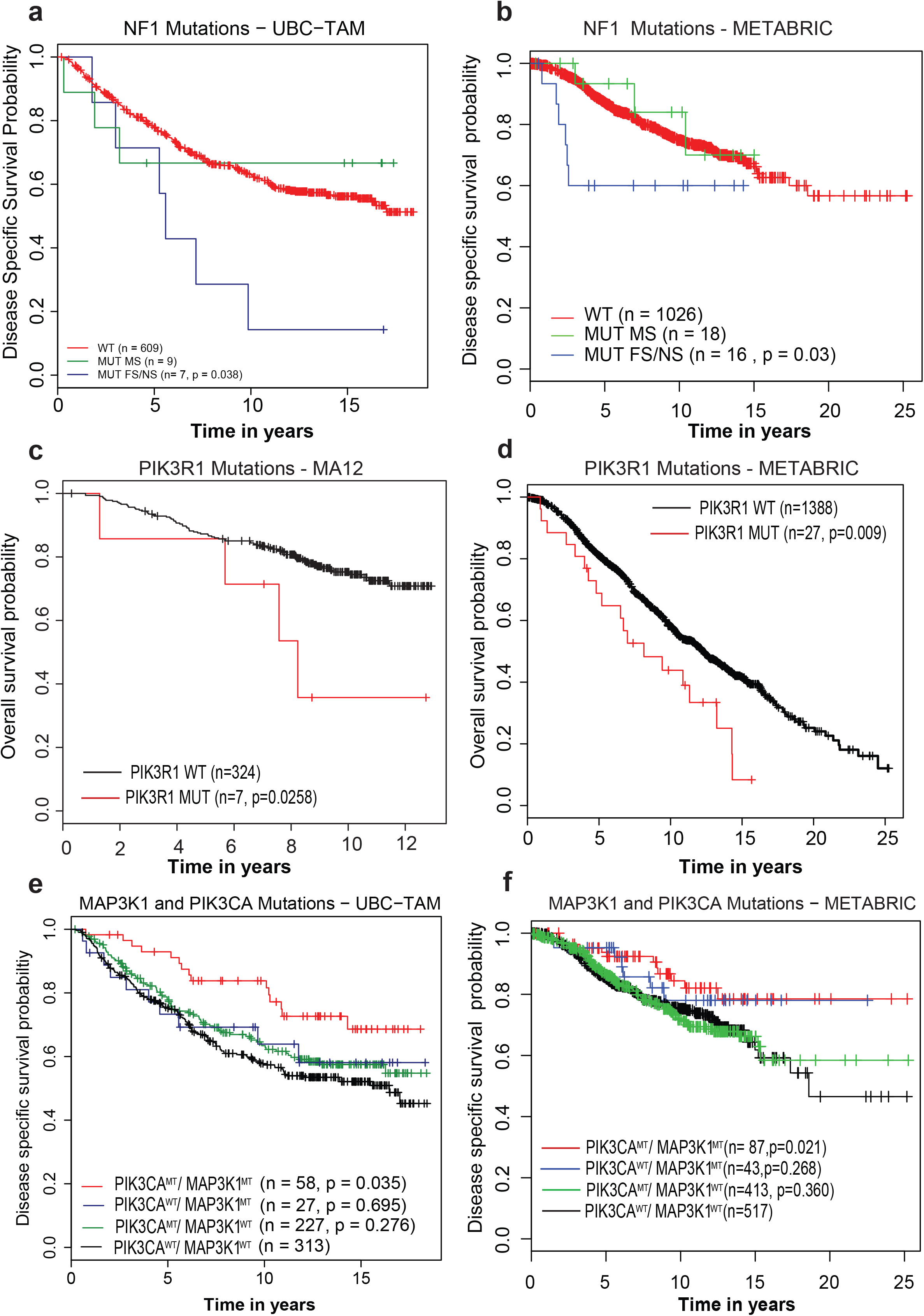
Kaplan-Meier plots. A-B) Kaplan-Meier graph showing the prognostic role of NF1 mutations, separated by variant type – Missense (MUT MS, green), Frameshift/Nonsense (MUT FS/NS, blue) in ER+ breast cancer patients from A) UBC-TAM and B) METABRIC cohort establishing the association between FS/NS mutations in NF1 with poor prognosis. C-D) Kaplan-Meier graph showing the prognostic role of PIK3R1 in C) MA12 and D) METABRIC ER+ breast cancer patients, categorized based on tumors with wildtype (WT, black) or mutated PIK3R1 non-silent mutations (MUT, red). E-F) Kaplan-Meier graph demonstrating co-occurrence of non-silent mutations in MAP3K1 and PIK3CA (red) in E) UBC-TAM and F) METABRIC associates with better survival when compared against tumors with mutations exclusively in MAP3K1 (blue) or PIK3CA (green) or wildtype for both MAP3K1 and PIK3CA (black). p, log rank (Mantel-Cox) test p-value.

### Prognostic interactions between PIK3CA and MAP3K1

Since PIK3CA and MAP3K1 mutations co-associate, the combined effect of non-silent mutations in these genes was examined. Patients with tumors exhibiting both genes mutated have a more favorable clinical course than either singly mutant cases or cases without either gene mutated. While the prognostic effects were strongest in the UBC-TAM series, this result was also reproduced in the METABRIC data (**Figure 7E-F**).

### Mutation Analyses for Uncommon Targetable Kinases

Of the 83 genes analyzed, at least 8 are directly targetable with small molecules or antibodies that are either FDA approved or in late-stage development (**Figure 8**). Pre-existing data on these mutations is summarized (Supplementary Data 7). PIK3CA is not further discussed here, since the mutation spectrum is well-described and large therapeutic studies are already underway. A total of 23 patients with breast cancer with ErbB2 activating mutations were identified. An examination of their locations revealed that ErbB2 mutations were, as expected, clustered in 2 major domains, with 2 of 23 having extracellular domain mutations at residue 310 and 21 of 23 having kinase domain mutations between residues 755-842^8, 15^. To further investigate the preliminary finding of an adverse prognostic effect for ErbB2 mutation in the MA12 series, an examination of the METABRIC data indicated that known activating mutations in ErbB2 were associated with a near significant adverse effect (HR=1.71, P=0.075) (Supplementary Figure 8). For ERBB3, 2 known-activating mutations were identified (V104L and E928A)^16^. The DDR1 kinase domain mutation, R776W, is possibly homologous to EGFR hot spot mutation L858R, but the remaining DDR1 variants are of unknown significance. For the mutations in JAK1, 3 of 12 are loss of function mutations (frame shift or non-sense) and the S816* mutation has been reported in a lung adenocarcinoma sequencing data set^17^. The loss of function mutations in JAK1 have been shown to associate with immune therapy resistance^18, 19^. A few mutations identified in ERBB4, MET, and PDGFRA have been previously reported but those reported here have not been functionally tested.

**Figure 8.**
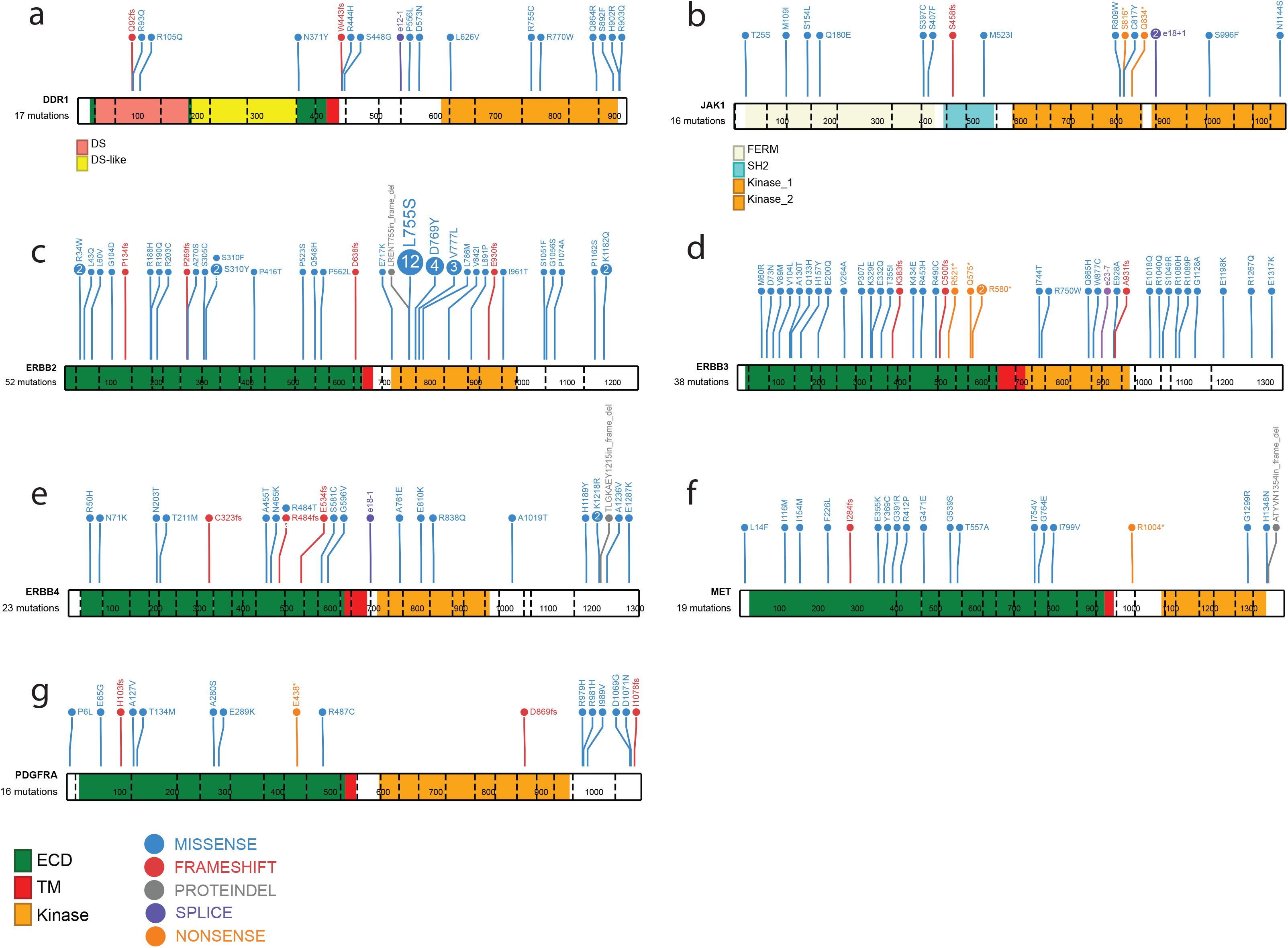
Mutation profiles for selected genes. Mutation frequency plots illustrate all non-silent mutations (TAM, POLAR, and MA12; n=1259) for representative transcripts for several kinase genes of interest. The domains belonging to A) DDR1 (RefSeq ID: NM_013994) and B) JAK1 (NM_002227) are indicated below the schematic diagram of each gene. The ECD (extracellular domain), TM (transmembrane domain), and kinase domain are depicted as green, red, and orange bars respectively for C) ERBB2 (NM_004448), D) ERBB3 (NM_001982), E) ERBB4 (NM_005235), F) MET (NM_000245), and G) PDGFRA (NM_006206). The variant counts across the three datasets for each gene are provided below the gene’s name. Note, in the mapping from Ensembl (**Supplementary Data 3**) to RefSeq annotations (required for use of ProteinPaint tool) a small number of variants annotations may have changed or been lost, despite selecting the most similar representative transcript possible.

## Discussion

The landscape of recurrently mutated genes in ER+ breast cancer observed in this study is consistent with reports where matched germline samples were available, indicating that our variant filters were effective for somatic mutation detection in a research setting. Overall, mutation rates were higher in our cohort (e.g., for *PIK3CA, MLL3, MAP3K1*) than the TCGA cohort, but were also lower for a few specific genes (e.g., *TP53* and *GATA3*). Due to higher sequencing data coverage of recurrently mutated target genes than TCGA and the use of a different hybrid capture reagent, we were likely able to detect mutations that were missed with lower-depth exome or whole genome sequencing data. It is also possible that in some instances we overestimated somatic mutation rates, due to the lack of matched normal samples and imperfections in our germline polymorphism filtering. In particular, a significant number of BRCA1 and BRCA2 mutations are likely *de novo* germline mutations that we would not be able to easily distinguish from somatic mutations. Of the 117 non-silent BRCA1/2 mutations observed (from 110/1128 patients; 7 patients had two hits) 74 were observed at a VAF greater than 40% and 31 were greater 60%. Variants with VAFs this high are less likely to be somatic given the general expectation of impure tumor samples and heterozygous mutations. Indeed, the VAFs for BRCA1/2 non-silent mutations (mean=44.8%) were significantly higher than for other genes (mean=36.4%, p = 6.96e-06). There were 8 known pathogenic (ENIGMA expert reviewed) mutations according to a search of the BRCA Exchange database (<u>http://brcaexchange.org</u>, Nov 12, 2017) and another 37 likely pathogenic (FS/NS) mutations. Of the remaining, 4 were known benign according to expert review (ENIGMA), and 8 benign, 15 likely benign and 45 variants of unknown significance according to all public sources.

The discovery of a novel recurrent *CBFB* (core binding factor subunit beta) splice site mutation in this cohort illustrates a limitation of exome capture reagents. The affected bases in exon 2 of CBFB display reduced sequence coverage, possibly due to high GC content, in the breast TCGA exome dataset (Supplementary Figures 9-10). This site was mutated in at least 1.5% of ER+ breast cancers sequenced, bringing the overall rate of CBFB mutations to nearly 6%, which should drive further investigation of this gene in ER+ breast cancer pathogenesis. *CBFB* functions as a subunit in a heterodimeric core binding transcription factor that interacts with *RUNX1*^20^. Consistent with this model, *CBFB* mutants were mutually exclusive from *RUNX1* mutants in this cohort with only a single sample harboring non-silent mutations in both *CBFB* and *RUNX1*.

The UBC-TAM and MA12 studies revealed different lists of potentially prognostic mutations. Prognostic effects are likely to be strongly affected by the use of systemic therapy as well as by patient age at diagnosis. The UBC-TAM series is the simplest study to interpret from a drug resistance perspective since the only systemic therapy was tamoxifen. Thus, the consistent adverse effect of NF1 FS/NS mutation on prognosis is intriguing as this result is consistent with results from an *in vitro* screen for tamoxifen resistance^21^. Understanding why only FS/NS mutations predict poor outcome, rather than missense or other non-silent mutations, will require further investigation. In contrast, PIK3R1 mutation emerged as a consistent poor prognosis mutation from the MA12 analysis, with validation in METABRIC. The proposed favorable prognostic effects of PIK3CA mutation were observed in the UBC-TAM series, but were not found to be independent of stage and grade, and PTEN mutations were neutral.

According to our validation results, NF1, PIK3R1, MAP3K1, PIK3CA and TP53 are likely to be prognostic drivers. In postmenopausal women treated with adjuvant endocrine therapy, DDR1, PRKDC and XBP1 should be further studied and of these DDR1 is the strongest candidate because it was significant despite strict false discovery correction. DDR1 is a collagen-binding receptor expressed in epithelial cells that stabilizes E-cadherin–mediated intracellular adhesion^22^. *DDR1* mutations also occur in endometrial cancer^23^, acute leukemia^24^ and lung cancer^25^. Loss of DDR1 (DDR1-null mice) produces hyper-proliferation and abnormal branching of mammary ducts, suggesting DDR1 is a breast tumor suppressor^26^. The relationship between truncating mutations in NF1 and poor outcome is consistent with an siRNA screen for genes whose loss generates tamoxifen resistance^21^. Mutations in PRKDC will potentially produce a defective ATM response/low ATM levels^27^ which is interesting in the context of the finding herein that ATM mutations are a potential luminal B driver gene. The significance of a defective ATM pathway as a cause of endocrine resistance is highlighted by the recent finding that dysregulation of the MutL complex (MLH1, PMS1 and PMS2) causes failure of ATM/CHK2-based negative regulation of CDK4/6^28^. Prognostic candidate mutations revealed by the MA12 analysis were different from the UBC TAM series, likely reflecting the different patient profiles and adjuvant treatments illustrated in **Figure 2**. The prognostic effects of mutations ERBB2, ERBB4, JAK1, LTK, MAP3K4, MET, PDGFRA, RB1, RELN, TGFB2, all await further study with even larger sample sizes.

In conclusion, we have successfully utilized clinically well-annotated, uniformly treated patient samples using DNA from archival material greater than 20 years old that lacks a matched normal to explore the prognostic effects encoded by the mutational landscape of ER+ breast cancer. We were able to confirm our prospective hypothesis that MAP3K1 is associated with indolent disease and TP53 with adverse outcomes. We also associated NF1 FS/NS mutations with strong adverse effects on prognosis. Similarly, PIK3R1 mutations were associated with an adverse prognosis in contrast to PIK3CA mutation. This suggests somatic mutations in these two physically interacting gene products are not biologically equivalent with respect to PI3 kinase pathway activation and resistance effects. The possibility that the long tail of low frequency mutation events in luminal type breast cancer may harbor multiple molecular explanations for poor outcomes is an important finding that should spur collaborative efforts to thoroughly screen thousands of properly annotated cases. Only after these iterative efforts of proposing and confirming candidates will a clinically useful and comprehensive somatic mutation-based classification of ER+ breast cancer emerge. In the meantime, functional studies should be pursued to understand the biological effects of somatic mutations, prioritizing these studies according to whether the mutations are driving an adverse prognostic effect.

## Methods

For the UBC-TAM series, an institutional review board approved study was based on formalin-fixed paraffin embedded (FFPE) primary tumor blocks from 947 female patients diagnosed with estrogen receptor positive invasive breast cancer in the province of British Columbia in Canada between 1986 and 1992^6, 29–31^. The sample flow and analysis are provided in a REMARK summary (**Figure 3A**). DNA was isolated from tumor-rich regions using the Qiagen blood and tissue kit, which yielded sufficient DNA in 645 samples, of which 625 met all study criteria and had sufficient sequence coverage. Similarly, approved studies provided 194 and 454 HR+ patient samples for the POLAR and MA12 (**Figure 4A**) cohorts. A total of 175 POLAR and 459 (328 HR+) MA12 samples yielded sufficient DNA and had sufficient sequence coverage for analysis. Detailed descriptions of the patient data sets are provided in Supplementary Table 3. A meta-analysis of six existing published large-scale breast cancer sequencing studies^1–3, 5, 32, 33^ was performed to identify genes with recurrent coding region somatic mutations in breast cancer (Supplementary Data 1). Additional drug targets^34^ and genes with relevance to breast cancer from targeted sequencing^35^, copy-number studies^13^ or knowledge relating to somatic or germline mutations (e.g., *BRCA1, BRCA2, ERBB2, ESR1* and *PRLR*) were also included. This resulted in a final list of 83 breast-cancer-related genes (Supplementary Table 1). These genes were targeted comprehensively with 3,029 complementary probes for hybridization-based enrichment (Supplementary Data 8). Sequencing libraries were constructed, hybridized with capture probes, multiplexed and run on a single flow cell with up to 96 samples per pool per lane yielding approximately 375 Mb of DNA sequence per sample from an Illumina HiSeq paired end 2 X 100bp (TAM) or 2 X 125bp (POLAR, MA12) sequencing run following manufacturer’s protocols.

Variant calling was performed with the Genome Modeling System as previously described^36^. Specifically, sequence data were aligned to reference sequence build GRCh37 using BWA^37^ and de-duplicated with Picard. SNVs and indels were detected using the union of samtools^38^ and VarScan2^5^ and annotated using Ensembl version 70. Variants were restricted to the coding regions of targeted genes and filtered for false positives and germline polymorphisms against a database of nearly 70,000 unmatched normals from the ExAC consortium, 1000 Genomes^39^, NHLBI exomes^40^ and TCGA data sets^3, 41^. A binomial probability model was then applied to the variants using VAF and total coverage to determine a log-likelihood ratio of being a somatic variant as previously described^42^ (See Supplementary Methods). After filtering, all remaining variants were manually reviewed. To ensure that variants of known clinical relevance were not missed by automated variant calling approaches, a knowledge-based variant calling strategy was performed focused on the mutations in the Database of Curated Mutations^43^.

Patient groups were defined by mutation status or truncating mutation status for each gene. Fisher’s exact and Chi-squared tests were used for hotspot analysis, mutual exclusivity or co-occurrence, and other categorical clinical statistics (e.g., mutation status vs. intrinsic subtype) as appropriate. Univariate Kaplan-Meier and Cox survival analyses were performed for breast-cancer-specific survival (BCSS), relapse free survival (RFS), or overall survival (OS) with non-silent or truncating mutation status as a factor. Significant survival differences between the groups were determined by log rank (Mantel-Cox) test. The Benjamini-Hochberg method was performed for multiple testing corrections to report the false discovery rate adjusted p-value (q-value). A multivariate Cox proportional hazard model was fitted to BCSS and RFS separately on gene mutation status, node status, grade and tumor size and adjusted hazard ratios were calculated with Wald test p-values. All statistical analyses were performed in the R statistical programming language with core, ‘survival’ and ‘multtest’ libraries. Genomic visualizations were created with ProteinPaint^44^ and GenVisR^45^.

## Acknowledgements

Research reported in this publication was primarily supported by Susan G. Komen Promise grant (PG12220321 to MJE), a Cancer Prevention and Research Institute of Texas (CPRIT) Recruitment of Established Investigators award (RR140033 to MJE). Dr. Ellis is a McNair Medical Institute Investigator and a Susan G. Komen Scholar. SMK was supported by a Komen CCR award (CCR16380599). The MA12 analysis was supported by research grants from Canadian Cancer Society Research Institute to the NCIC Clinical Trials Group (021039 and 015469). OLG was supported by the National Cancer Institute (NIH NCI K22CA188163 and NIH NCI U01CA209936).

## Contributions

O.L.G., N.C.S., T.O.N., M.J.E., E.R.M. designed the experiments; M.G., J. K., C.A.M., K.K., J.H., B.J.A., Z.L.S., K.C. R.K. C.F., L.C., J.E.S., S.D., V.M., D.E.L., R.S.F., S.L., R.K.W. generated the sequencing data, T.O.N., B.Y., M.D. S.L., and D.V. orchestrated the sample pipeline., M.A., O.L.G. and N.C.S., prepared the figures and tables. M.A., J.L., and D.T. provided statistical analysis. S.M.K., R.B., and E.C.C. provided functional annotations., T.O.N provided pathology analysis. M.J.E., N.C.S., M.A., and O.L.G. wrote the manuscript. E.R.M., T.O.N., M.D., critically read and commented on the manuscript.

